# Reduced Lateralization of Multiple Functional Brain Networks in Autistic Males

**DOI:** 10.1101/2023.12.15.571928

**Authors:** Madeline Peterson, Molly B. D. Prigge, Dorothea L. Floris, Erin D. Bigler, Brandon Zielinski, Jace B. King, Nicholas Lange, Andrew L. Alexander, Janet E. Lainhart, Jared A. Nielsen

**Affiliations:** Department of Psychology, Brigham Young University, Provo, UT, 84602, USA; Department of Radiology and Imaging Sciences, University of Utah, Salt Lake City, UT, 84108, USA; Waisman Center, University of Wisconsin-Madison, Madison, WI, 53705, USA; Methods of Plasticity Research, Department of Psychology, University of Zurich, Zurich, Switzerland; Department of Cognitive Neuroscience, Donders Institute for Brain, Cognition and Behaviour, Radboud University Nijmegen Medical Center, Nijmegen, The Netherlands; Neuroscience Center, Brigham Young University, Provo, UT, 84604, USA; Department of Neurology, University of Utah, Salt Lake City, UT, 84108, USA; Department of Neurology, University of California-Davis, Davis, CA, USA; Department of Pediatrics, University of Utah, Salt Lake City, UT, 84108, USA; Division of Pediatric Neurology, Departments of Pediatrics, Neurology, and Neuroscience, College of Medicine, University of Florida, FL, 32610, United States; Department of Psychiatry, Harvard Medical School, Boston, MA, 02115, USA; Department of Psychiatry, University of Wisconsin-Madison, Madison, WI, 53719, USA; Department of Medical Physics, University of Wisconsin-Madison, Madison, WI, 53705, USA

**Keywords:** autism, autism spectrum disorder, ASD, asymmetry, brain networks, fMRI, lateralization, network lateralization, neuroimaging, neurodevelopmental conditions

## Abstract

**Background:** Autism spectrum disorder has been linked to a variety of organizational and developmental deviations in the brain. One such organizational difference involves hemispheric lateralization, which may be localized to language-relevant regions of the brain or distributed more broadly.

**Methods:** In the present study, we estimated brain hemispheric lateralization in autism based on each participant’s unique functional neuroanatomy rather than relying on group-averaged data. Additionally, we explored potential relationships between the lateralization of the language network and behavioral phenotypes including verbal ability, language delay, and autism symptom severity. We hypothesized that differences in hemispheric asymmetries in autism would be limited to the language network, with the alternative hypothesis of pervasive differences in lateralization. We tested this and other hypotheses by employing a cross-sectional dataset of 118 individuals (48 autistic, 70 neurotypical). Using resting-state fMRI, we generated individual network parcellations and estimated network asymmetries using a surface area-based approach. A series of multiple regressions were then used to compare network asymmetries for eight significantly lateralized networks between groups.

**Results:** We found significant group differences in lateralization for the left-lateralized Language (d = −0.89), right-lateralized Salience/Ventral Attention-A (d = 0.55), and right-lateralized Control-B (d = 0.51) networks, with the direction of these group differences indicating less asymmetry in autistic individuals. These differences were robust across different datasets from the same participants. Furthermore, we found that language delay stratified language lateralization, with the greatest group differences in language lateralization occurring between autistic individuals with language delay and neurotypical individuals.

**Limitations:** The generalizability of our findings is restricted due to the male-only sample and greater representation of individuals with high verbal and cognitive performance.

**Conclusions:** These findings evidence a complex pattern of functional lateralization differences in autism, extending beyond the Language network to the Salience/Ventral Attention-A and Control-B networks, yet not encompassing all networks, indicating a selective divergence rather than a pervasive one. Furthermore, a differential relationship was identified between Language network lateralization and specific symptom profiles (namely, language delay) of autism.

## Background

Autism spectrum disorder (ASD) is a heterogenous neurodevelopmental condition characterized by challenges in social communication and the presence of restricted repetitive behaviors (Diagnostic and Statistical Manual-5; 1). As a neurodevelopmental condition, ASD is linked to atypical timelines of social, cognitive, and physiological development. A pivotal question in the study of autism revolves around the role that alterations in brain asymmetries may play in its development.

Hemispheric specialization, a key principle of brain organization and design, refers to the phenomenon whereby specific cognitive functions are predominantly localized in one hemisphere over the other. This specialization is akin to a division of labor within the brain, with each hemisphere assuming distinct yet not exclusive cognitive responsibilities. In essence, it is as if the brain has a dominant hand for certain types of cognitive operations, such as emotion responsiveness, visuospatial attention, conscious problem solving, and language processing, among others (2). The near universality of these functional asymmetries in the human brain raises intriguing questions regarding their behavioral and cognitive purposes. It has been hypothesized that the emergence of lateralized cognitive functions may have been a crucial adaptation that allowed humans to excel in various aspects of life, including improved mobility, more astute resource-seeking behavior, and more effective defense against predators (3). The implications for brain function are also intriguing, and it is thought that functional asymmetries reflect a dynamic trade-off between decreases in redundancy (4), processing speed (5), and interhemispheric conflict in function initiation (6,7), and the loss of system redundancies and inter-hemispheric connections.

When considering neurodevelopmental conditions such as ASD, the consequences of atypical hemispheric lateralization or a lack thereof become particularly relevant. Language laterality in particular is of interest to autism researchers, since the diagnosis includes a number of language-related features. Consequently, the field of autism research has a long history of investigating lateralization, employing a variety of research methods. For instance, a dichotic listening task paradigm identified a reversal or reduction of lateralization for speech in ASD (8). Subsequent electroencephalography studies arrived at a similar conclusion (9–12). However, despite evidence from these and additional studies, it is unknown if differences in hemispheric lateralization in autism are localized to language-relevant regions, as posited in the left hemisphere dysfunction theory of autism (13), or if they are more pervasive.

Current evidence surrounding this lateralization debate is inconsistent, with findings for generally increased activity in the right hemisphere in autism (14–18), generally decreased activity in the left hemisphere (19–21), both increased activity in the right hemisphere and decreased activity in the left hemisphere (22–24), and generally decreased connectivity across both hemispheres (25). Conversely, recent evidence for specific differences in lateralization for regions involved in language processing in autism is compelling. For example, in a functional connectivity-based study, a reduction in left lateralization was observed for several connections involving left-lateralized hubs, particularly those related to language and the default network (26). This was examined once more by Jouravlev et al. (27) with a functional language task on an individual level. Within the language network, autistic participants showed less lateralized responses due to greater right hemisphere activity (27). Interestingly, there was no strong difference in lateralization for the theory of mind and multiple demand networks between autistic and neurotypical (NT) participants, suggesting that differences in lateralization are constrained to language regions (27).

Adding another layer to this debate is the potential role that language delay might play in stratifying differences in lateralization in autism. Using normative modeling, one team found that language delay explained the most variance in extreme rightward deviations of laterality in autism (28). This is a promising direction, as it appears language delay is capable of parsing the heterogeneity of atypical lateralization patterns in autism. Furthermore, this result points to the behavioral relevance of atypical lateralization patterns to language development in autism. However, it is unclear as to if atypical lateralization in language regions specifically or global alterations of lateralization are contributing to the observed language deficits (29).

The aim of the present study is to address this ongoing debate regarding the specificity of atypical lateralization patterns to language-relevant regions in autism. This was undertaken by approaching both brain network parcellations and network lateralization from an individual level. The use of these individualized elements is non-trivial, since functional networks vary more by stable group and individual factors than cognitive or daily variation (30). Furthermore, group averaging can obscure individual differences and blur functional and anatomical details—details which are potentially clinically useful (31,32). Thus, through the use of this individual approach, we are better positioned to capture idiosyncratic functional and anatomical details relevant to network lateralization.

The present study explored following hypotheses. First, it was hypothesized that autistic individuals would show reduced hemispheric lateralization only in areas associated with language compared with neurotypical individuals. Second, we examined the relationships between language lateralization and three behavioral phenotypes: verbal ability, autism symptom severity, and language delay. More specifically, we hypothesized a positive relationship between language lateralization and verbal ability (as previously described by 33), and a negative relationship between language lateralization and autism symptom severity. Finally, we hypothesized that language delay would stratify language lateralization, with the greatest expected differences in lateralization to occur between autistic individuals with language delay and neurotypical individuals.

## Methods

### Participants

A previously collected dataset was used, and further information on participant recruitment and diagnosis can be found elsewhere (34–37). All data were obtained with assent and informed consent according to the University of Utah’s Institutional Review Board. Participants underwent two 15-minute resting-state multi-echo functional magnetic resonance imaging (fMRI) scans and were instructed to simply rest with their eyes open while letting their thoughts wander (38). A total of *N* = 89 ASD and *N* = 108 NT participants had fMRI data. Exclusion criteria included participants without age data, participants without handedness data (Edinburgh Handedness Inventory; 39), participants older than 50 years, female participants, participants with less than 50% of volumes remaining after motion censoring, and participants with a mean framewise displacement greater than 0.2 mm and mean DVARS greater than 50. The exclusion criterion of age greater than 50 was selected due to the lack of matched controls for participants older than 50. Female participants were excluded from the analyses due to their limited representation (*N* = 3). A total of *N* = 48 ASD and *N* = 70 NT were included in the final analysis. In summary, ASD mean age was 27.22 years, range 14.67–46.42 years; NT mean age was 27.92 years, range 16.33–46.92 years; overall mean age was 27.63 years. Additional demographic information can be found in Table 1.

**Table 1.**
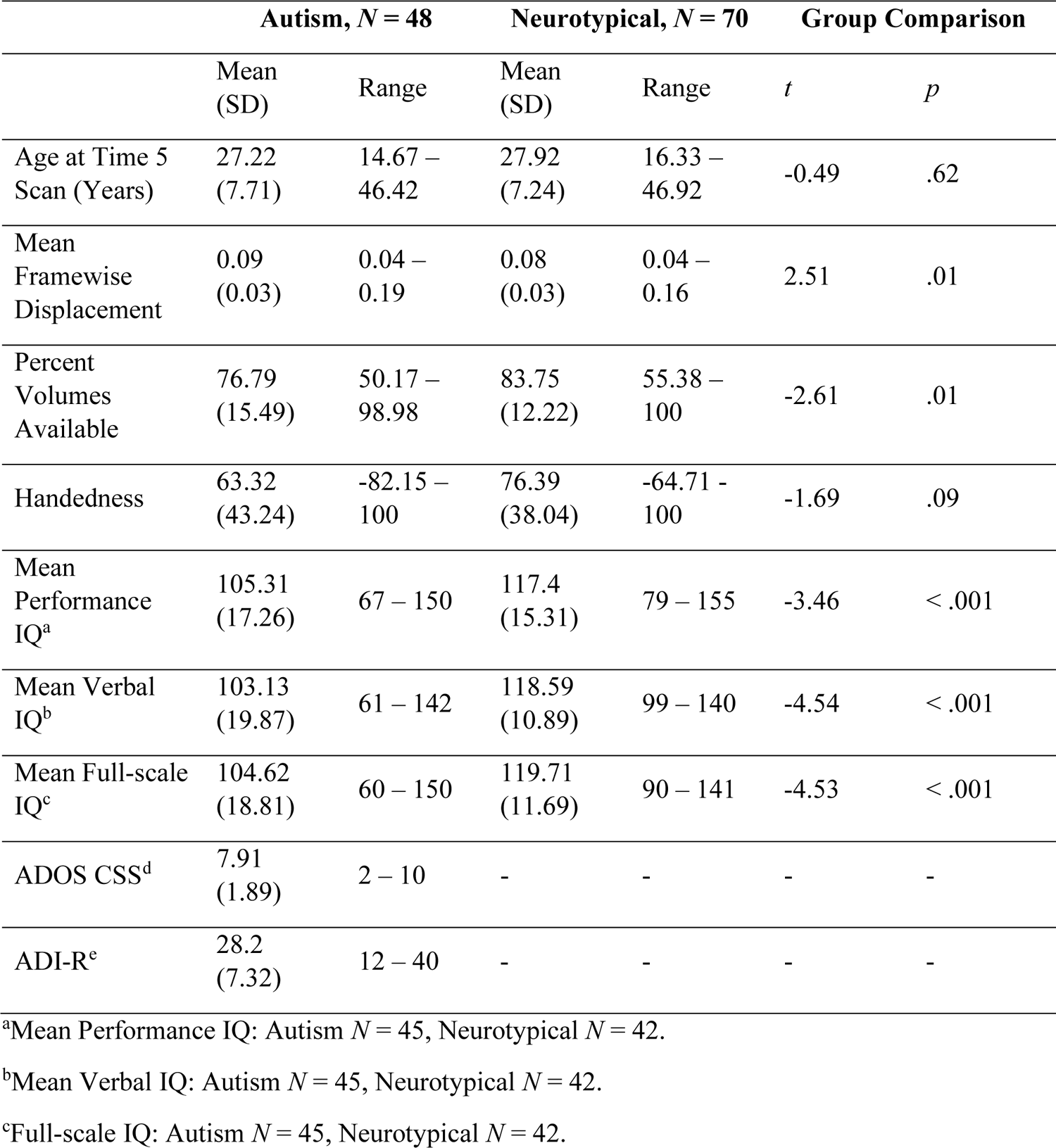
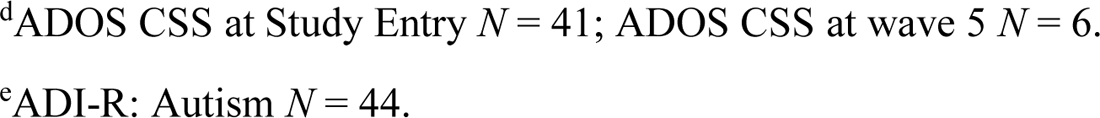
Demographics.

|  | Autism, *N* = 48 |  | Neurotypical, *N* = 70 |  | Group Comparison |  |
| --- | --- | --- | --- | --- | --- | --- |
|  | Mean<br>(SD) | Range | Mean<br>(SD) | Range | *t* | *p* |
| Age at Time 5<br>Scan (Years) | 27.22<br>(7.71) | 14.67 –<br>46.42 | 27.92<br>(7.24) | 16.33 –<br>46.92 | -0.49 | .62 |
| Mean<br>Framewise<br>Displacement | 0.09<br>(0.03) | 0.04 –<br>0.19 | 0.08<br>(0.03) | 0.04 –<br>0.16 | 2.51 | .01 |
| Percent<br>Volumes<br>Available | 76.79<br>(15.49) | 50.17 –<br>98.98 | 83.75<br>(12.22) | 55.38 –<br>100 | -2.61 | .01 |
| Handedness | 63.32<br>(43.24) | -82.15 –<br>100 | 76.39<br>(38.04) | -64.71 –<br>100 | -1.69 | .09 |
| Mean<br>Performance<br>IQ<sup>a</sup> | 105.31<br>(17.26) | 67 – 150 | 117.4<br>(15.31) | 79 – 155 | -3.46 | < .001 |
| Mean Verbal<br>IQ<sup>b</sup> | 103.13<br>(19.87) | 61 – 142 | 118.59<br>(10.89) | 99 – 140 | -4.54 | < .001 |
| Mean Full-scale<br>IQ<sup>c</sup> | 104.62<br>(18.81) | 60 – 150 | 119.71<br>(11.69) | 90 – 141 | -4.53 | < .001 |
| ADOS CSS<sup>d</sup> | 7.91<br>(1.89) | 2 – 10 | - | - | - | - |
| ADI-R<sup>e</sup> | 28.2<br>(7.32) | 12 – 40 | - | - | - | - |

<sup>a</sup>Mean Performance IQ: Autism *N* = 45, Neurotypical *N* = 42.
<sup>b</sup>Mean Verbal IQ: Autism *N* = 45, Neurotypical *N* = 42.
<sup>c</sup>Full-scale IQ: Autism *N* = 45, Neurotypical *N* = 42.
<sup>d</sup>ADOS CSS at Study Entry $N = 41$ ; ADOS CSS at wave 5 $N = 6$ .
<sup>e</sup>ADI-R: Autism $N = 44$ .

Autistic participants and neurotypical participants did not significantly differ in mean age (*t*(96.87) = −0.49, *p* = .62) or handedness (*t*(92.38) = −1.69, *p* = .09). However, the two groups did differ in data quality (*t*(93.81) = 2.51, *p* = .01) and quantity (*t*(85.13) = −2.61, *p* = .01).

Furthermore, there was a significant difference between groups on available intelligence quotient (IQ) measures (*p* < .001). Details regarding IQ measures in this dataset have been previously reported (35,37). Using full-scale IQ score of 79 or lower as the criterion for low verbal and cognitive performance (40), there were three autistic participants who met this criterion. Additionally, 93.33% of the autistic participants had high verbal and cognitive performance and 100% of the NT sample had high verbal and cognitive performance.

Table 1 also presents the Autism Diagnostic Observation Schedule (ADOS) calibrated severity scores (CSS) at entry. The ADOS was administered by trained clinicians or research-reliable senior study staff as detailed previously (35,37). The ADOS CSS scores were then calculated based on ADOS module and participant age (41). A few participants had ADOS CSS scores derived more recently (*N* = 6). The ASD diagnosis of these select participants was confirmed prior to study enrollment, so the ADOS was not administered at study entry to these participants. Autism Diagnostic Interview-Revised (ADI-R) scores are also reported, and these scores act as a summary of autism severity during childhood.

Characteristics of autistic participants with and without language delay can be found in Table 2. In accordance with prior work (28,42), language delay was operationalized as having the onset of first words later than 24 months and/or having onset of first phrases later than 33 months as assessed via the ADI (not the ADI-R). These ADI items were available for 45/48 autistic participants, of which 29 met the threshold for language delay.

**Table 2.**
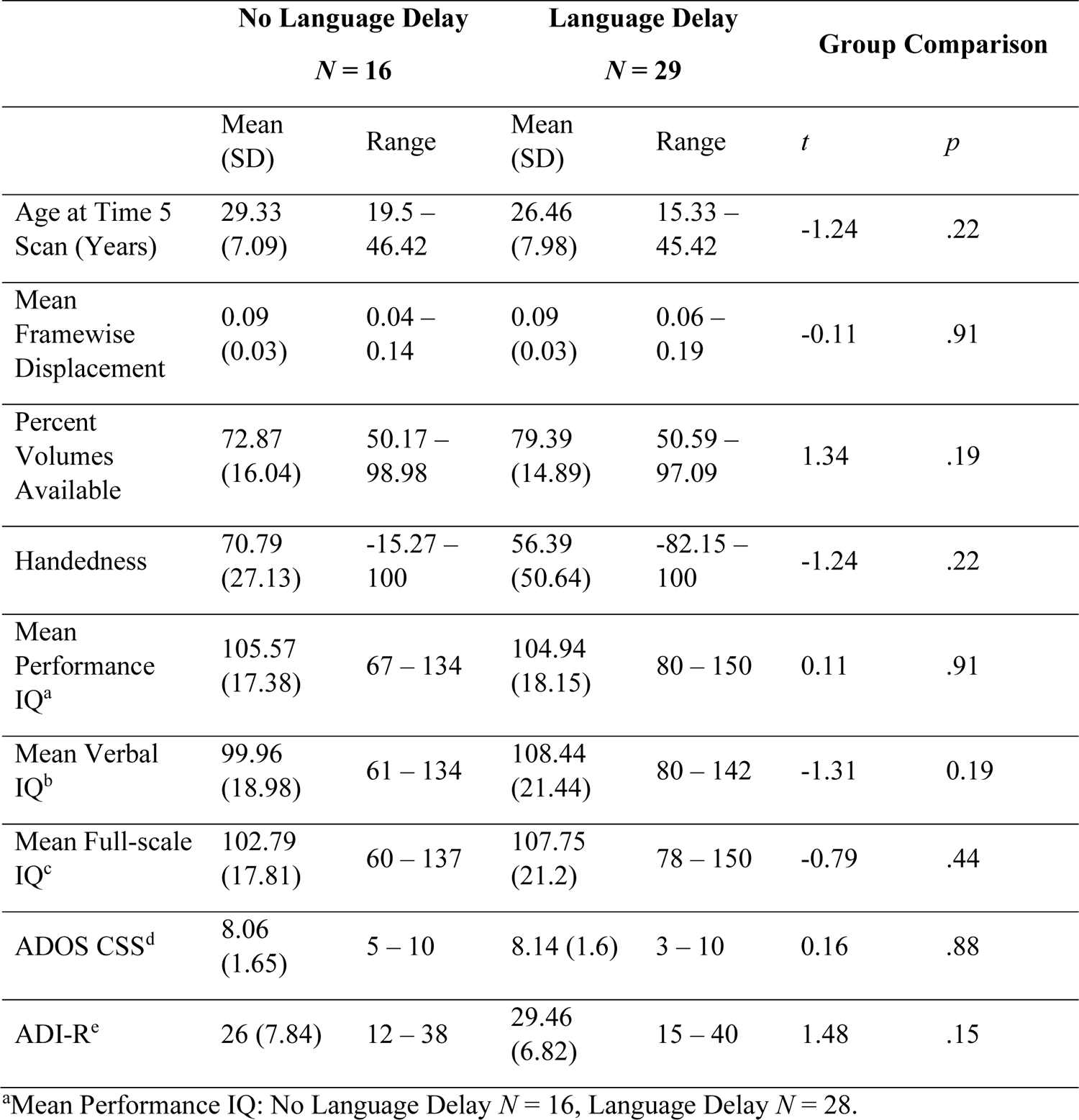
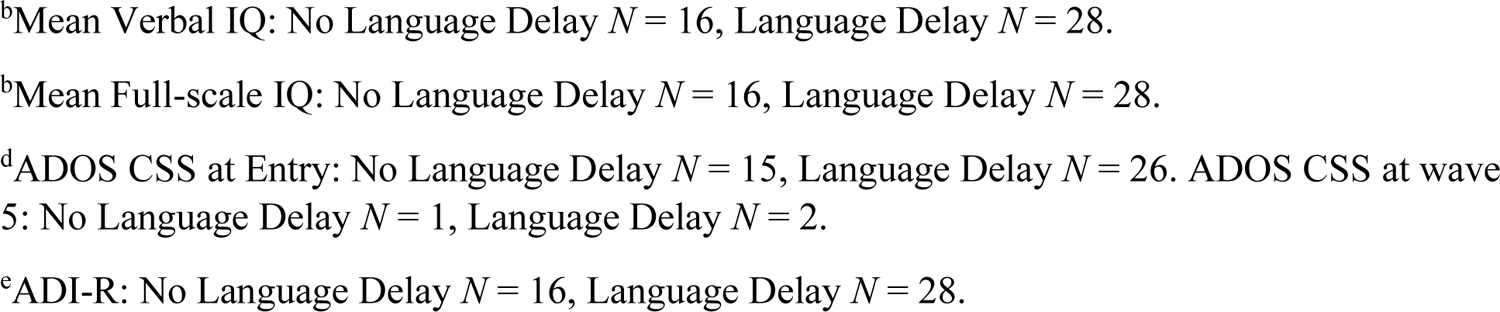
Language Delay Demographics.

|  | No Language Delay |  | Language Delay |  | Group Comparison |  |
| --- | --- | --- | --- | --- | --- | --- |
|  | *N* = 16 |  | *N* = 29 |  | *t* | *p* |
|  | Mean (SD) | Range | Mean (SD) | Range |  |  |
| Age at Time 5 Scan (Years) | 29.33 (7.09) | 19.5 – 46.42 | 26.46 (7.98) | 15.33 – 45.42 | -1.24 | .22 |
| Mean Framewise Displacement | 0.09 (0.03) | 0.04 – 0.14 | 0.09 (0.03) | 0.06 – 0.19 | -0.11 | .91 |
| Percent Volumes Available | 72.87 (16.04) | 50.17 – 98.98 | 79.39 (14.89) | 50.59 – 97.09 | 1.34 | .19 |
| Handedness | 70.79 (27.13) | -15.27 – 100 | 56.39 (50.64) | -82.15 – 100 | -1.24 | .22 |
| Mean Performance IQ<sup>a</sup> | 105.57 (17.38) | 67 – 134 | 104.94 (18.15) | 80 – 150 | 0.11 | .91 |
| Mean Verbal IQ<sup>b</sup> | 99.96 (18.98) | 61 – 134 | 108.44 (21.44) | 80 – 142 | -1.31 | 0.19 |
| Mean Full-scale IQ<sup>c</sup> | 102.79 (17.81) | 60 – 137 | 107.75 (21.2) | 78 – 150 | -0.79 | .44 |
| ADOS CSS<sup>d</sup> | 8.06 (1.65) | 5 – 10 | 8.14 (1.6) | 3 – 10 | 0.16 | .88 |
| ADI-R<sup>e</sup> | 26 (7.84) | 12 – 38 | 29.46 (6.82) | 15 – 40 | 1.48 | .15 |

<sup>a</sup>Mean Performance IQ: No Language Delay *N* = 16, Language Delay *N* = 28.
<sup>b</sup>Mean Verbal IQ: No Language Delay $N = 16$ , Language Delay $N = 28$ .
<sup>b</sup>Mean Full-scale IQ: No Language Delay $N = 16$ , Language Delay $N = 28$ .
<sup>d</sup>ADOS CSS at Entry: No Language Delay $N = 15$ , Language Delay $N = 26$ . ADOS CSS at wave 5: No Language Delay $N = 1$ , Language Delay $N = 2$ .
<sup>e</sup>ADI-R: No Language Delay $N = 16$ , Language Delay $N = 28$ .

### MRI Acquisition Parameters

MRI data were acquired at the Utah Center for Advanced Imaging Research using a Siemens Prisma 3T MRI scanner (80 mT/m gradients) with the vendor’s 64-channel head coil (see (38); Siemens, Erlangen, Germany). Structural images were acquired with a Magnetization Prepare 2 Rapid Acquisition Gradient Echoes (MP2RAGE) sequence with isotropic 1-mm resolution (Repetition Time (TR) = 5000 milliseconds, Echo Time (TE) = 2.91 milliseconds, and inversion time = 700 milliseconds). Resting-state functional images were acquired with a multiband, multi-echo, echo-planar sequence (TR = 1553 milliseconds; flip angle = 65°; in-plane acceleration factor = 2; fields of view = 208 mm; 72 axial slices; resolution = 2.0 mm isotropic; multiband acceleration factor = 4; partial Fourier = 6/8; bandwidth = 1850 Hz; 3 echoes with TEs of 12.4 milliseconds, 34.28 milliseconds, and 56.16 milliseconds; and effective TE spacing = 22 milliseconds).

### fMRI Preprocessing

Preprocessing took place on raw Neuroimaging Informatics Technology Initiative (NIfTI) files for the resting-state fMRI runs using a pipeline developed by the Computational Brain Imaging Group (CBIG; 43,44). Briefly, preprocessing steps included surface reconstruction (using FreeSurfer 6.0.1; 45), removal of the first four frames (using FSL, or FMRIB Software Library; 46,47), multi-echo integration and denoising (using *tedana*; 48, structural and functional alignment using boundary-based registration (using FsFast; 49), linear regression using multiple nuisance regressors (using a combination of CBIG in-house scripts and the FSL MCFLIRT tool; 46), projection to FreeSurfer fsaverage6 surface space (using FreeSurfer’s *mri_vol2surf* function), and smoothing with a 6 mm full-width half-maximum kernel (using FreeSurfer’s *mri_surf2surf* function; 50). To take full advantage of the multi-echo echo planar image scans in this dataset, the parameters of the CBIG preprocessing pipeline included *tedana* (48). Multi-echo data are acquired by taking three or more images per volume at echo times spanning tens of milliseconds (51,52). This provides two specific benefits: 1) Echoes can be integrated into a single time-series with improved blood oxygen level dependent contrast and less susceptibility artifact via weighted averaging, and 2) the way in which signals decay across echoes can be used to inform denoising (53). Therefore, to take advantage of these properties, *tedana* creates a weighted sum of individual echoes and then denoises the data using a multi-echo independent component analysis-based denoising method (48). Additionally, as suggested by Kundu et al. (54), bandpass filtering was not included as a preprocessing step for the multi-echo data.

### Individual Network Parcellation

After the implementation of multi-echo preprocessing, network parcellations were computed using a multi-session hierarchical Bayesian modeling pipeline (43). This pipeline was implemented in MATLAB R2018b (55). In summary, the pipeline estimates group-level priors from a training dataset (37 Brain Genomics Superstruct Project subjects; 43,56) and applies those to estimate individual-specific parcellations. A *k* of 17 networks was selected for all subjects, following the 17-network solution found in Yeo et al. (57). A Hungarian matching algorithm was then used to match the clusters with the Yeo et al. (57) 17-network group parcellation.

### Network Surface Area Ratio

Following the generation of individual network parcellations, lateralization was estimated using the network surface area ratio (NSAR) calculated in Connectome Workbench *wb_command* v1.5.0 (58). This measure was previously examined for validity and reliability (59) and is calculated on an individual basis for each of 17 networks. NSAR values range from −1.0 to +1.0, with negative values indicating left hemisphere lateralization for a given network and positive values indicate right hemisphere lateralization. NSAR values closer to zero indicate less lateralization (e.g., hemispheric symmetry).

## Statistical Analysis

### Validation of the Neurotypical Group

Before formally testing the hypotheses, the laterality pattern of the NT group was first validated using a series of multiple regressions. Models consisted of NSAR values as the dependent variable with the covariates of mean-centered age, mean-centered mean framewise displacement, and handedness index score (39).

### Group Differences in Network Lateralization

A within-dataset replication was first performed using participants with two available resting-state runs (*N* = 97; ASD = 37, NT = 60). A demographics table for this subset of individuals is available in the Supplementary Materials (see Supplementary Table 1), as is a table describing data quality across the two available runs (see Supplementary Table 2). Using this subset of individuals, the first hypothesis regarding group differences in lateralization was tested using the first resting-state run (the Discovery dataset) and then the second resting-state run (the Replication dataset). To compare hemispheric lateralization between ASD and NT individuals, a series of multiple regressions were performed first within the Discovery dataset and then within the Replication dataset. Individual parcellations and lateralization values were calculated separately for the Discovery and Replication datasets. Models consisted of NSAR values as the dependent variable, group (ASD and NT) as the independent variable, and the following covariates: mean-centered age, mean-centered mean framewise displacement, and handedness. Multiple comparisons were addressed via Bonferroni correction. Any networks with group differences identified in the Discovery dataset were tested in the Replication dataset.

Following the hypothesis testing in the Discovery and Replication datasets, models were implemented in all of the participants, with individual parcellations and corresponding lateralization values derived from all available data (the Complete dataset). Note that the Complete dataset includes 21 participants with only one available scan, and that NSAR values for participants with two available scans were derived from a single individual parcellation created using both scans as input. Models consisted of NSAR values as the dependent variable, group (ASD and NT) as the independent variable, and the following covariates: mean-centered age, mean-centered mean framewise displacement, and handedness. Only networks with group differences identified in the Discovery or Replication datasets were tested in the Complete dataset, with a corresponding Bonferroni correction. Effect sizes (Cohen’s *d*) for any potential group differences were calculated on contrasts extracted from the corresponding multiple regression model (60). To provide additional rigor, sub-analyses using nearest neighbor matching between the ASD and NT groups on the basis of mean framewise displacement, percent volumes available, and full-scale IQ were implemented using the R package MatchIt (61).

### Network Lateralization and Behavioral Phenotypes

To address the second hypothesis and examine the relationship between language network lateralization and verbal IQ across ASD and NT individuals, a multiple regression was used within the Complete dataset. Covariates included mean-centered age, mean-centered mean framewise displacement, and handedness. A similar analysis including language lateralization as a predictor of autism symptom severity (measured via ADOS CSS scores) was also performed.

Lastly, the potential relationship between language delay and language lateralization in ASD was investigated within the Complete dataset. For these analyses, language lateralization measured via NSAR was the dependent variable while the predictor was group (NT, ASD Language Delay, and ASD No Language Delay), and covariates included mean-centered age, mean-centered mean framewise displacement, and handedness. All statistical analyses took place in R 4.2.0 (62).

## Results

### Validation of the Neurotypical Group

In order to validate the neurotypical group as a reference group for the group analysis, multiple regressions were used to identify significantly lateralized networks, and eight networks were identified as being lateralized: Visual-B, Language, Dorsal Attention-A, Salience/Ventral Attention-A, Control-B, Control-C, Default-C, and Limbic-B (see Supplementary Table 3 and Supplementary Figure S1). This result aligns with prior findings (59), validating the neurotypical group from the Complete dataset as a reference group.

### Group Differences in Network Lateralization

To test the hypothesis regarding group differences in lateralization, regression models were first implemented in Discovery and Replication datasets (composing a within-dataset replication) followed by the Complete dataset.

The first hypothesis regarding group differences in lateralization was examined using a series of multiple regressions. Adjusted for multiple comparisons, group differences in the Discovery dataset were found in the following networks: Language (*t*(92) = −3.18, *p*-adjusted = .02), Salience/Ventral Attention-A (*t*(92) = 3.82, *p*-adjusted = .002), and Control-B (*t*(92) = 3.06, *p*-adjusted = .02; see Figure 1 Panel B). Significant group differences in lateralization were identified for the Replication dataset for the Language (*t*(92) = −2.44, *p*-adjusted < .05) and Control-B (*t*(92) = 2.55, *p*-adjusted = .04) networks, but not for the Salience/Ventral Attention-A network (*t*(92) = 1.83, *p*-adjusted = .21; see Figure 1 Panel C). For a depiction of lateralization for all eight lateralized networks across the Discovery and Replication datasets, see Supplementary Figure S2.

**Figure 1.**
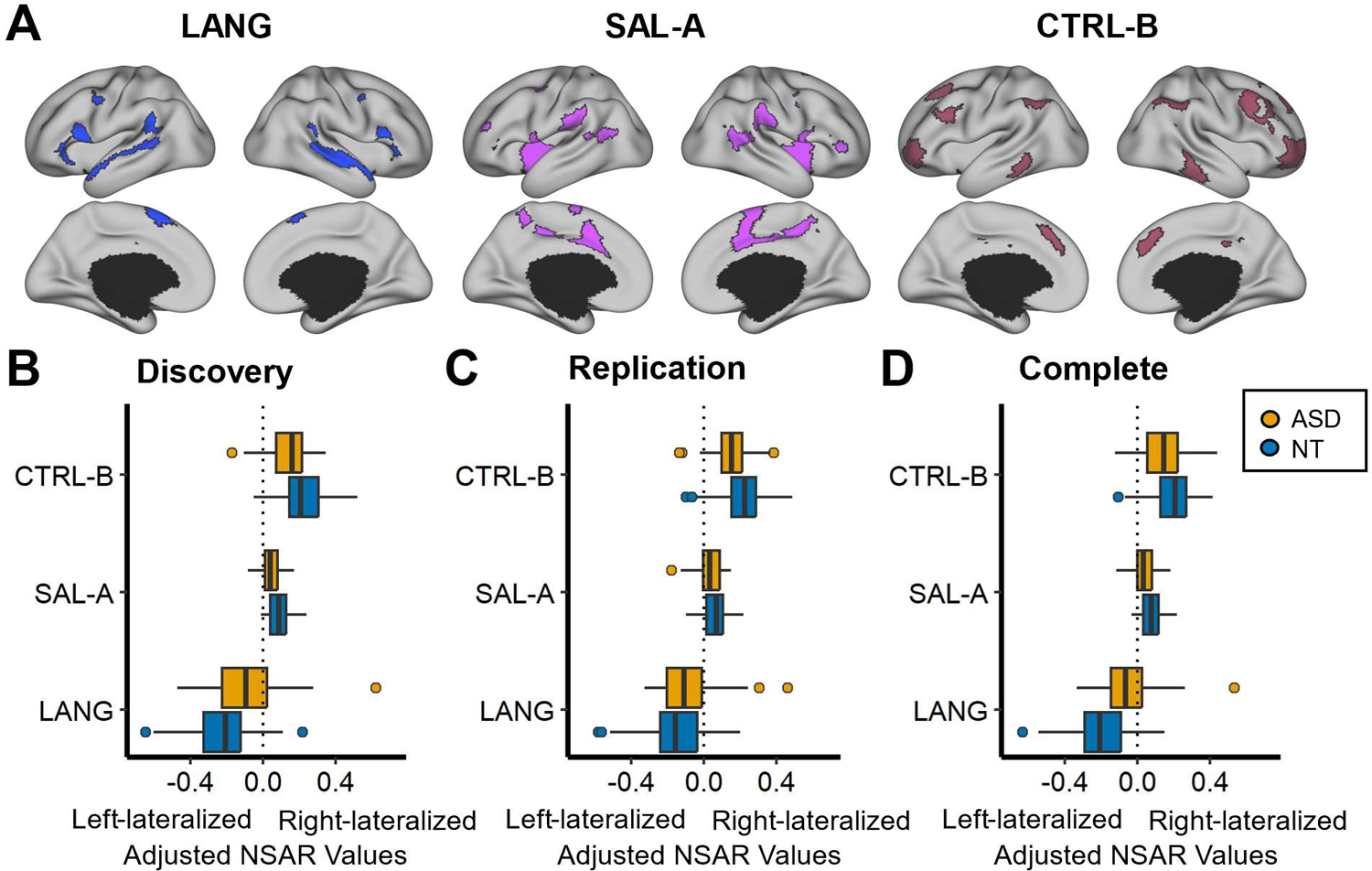
Group differences in network lateralization. Panel A depicts an individual parcellation from a neurotypical subject of three networks for which group differences in lateralization were identified. These networks include the Language (LANG), Salience/Ventral Attention-A (SAL-A), and Control-B (CTRL-B) networks. Panels B-D depict three networks on the y-axis and model-adjusted NSAR values on the x-axis, with negative values representing left hemisphere lateralization and positive values representing right hemisphere lateralization. NSAR values were adjusted by regressing out the effects of mean-centered age, mean-centered mean framewise displacement, and handedness using the following formula: NSAR_adjusted_ = NSAR_raw_ - [β_1_(mean-centered age_raw_ - mean of mean-centered age_raw_) + β_2_(mean-centered FD_raw_ - mean of mean-centered FD_raw_) + β_3_(group_raw_ - mean group_raw_) + β_4_(handedness_raw_ - mean handedness_raw_)]. NSAR adjustment occurred separately for each network and each group. A significant group effect on lateralization was found for three networks following Bonferroni correction in the Discovery dataset: Language (*t*(92) = −3.18, *p*-adjusted = .02), Salience/Ventral Attention-A (*t*(92) = 3.82, *p*-adjusted = .002), and Control-B (*t*(92) = 3.06, *p*-adjusted = .02). Significant group differences in lateralization for the Language (*t*(92) = −2.44, *p*-adjusted = .05) and Control-B (*t*(92) = 2.55, *p*-adjusted = .04) networks were replicated in the Replication dataset. In the Complete dataset, group differences in lateralization were identified for the Language (*t*(113) = −4.69, *p*-adjusted < .001), Salience/Ventral Attention-A (*t*(113) = 2.89, *p*-adjusted = .01), and Control-B (*t*(113) = 2.71, *p*-adjusted = .02) networks.

Next, multiple regressions were used to examine potential differences between the ASD and NT groups in lateralization in the Complete dataset for the three networks previously identified in the Discovery and Replication datasets. A significant group effect on lateralization was found for the three networks after Bonferroni correction: Language (*t*(113) = −4.69, *p*-adjusted < .001, d = −0.89), Salience/Ventral Attention-A (*t*(113) = 2.89, *p*-adjusted = .01, d = 0.55), and Control-B (*t*(113) = 2.71, *p*-adjusted = .02, d = 0.51; see Figure 1 Panel D). In order to understand which hemisphere was driving differences in lateralization, we examined network surface areas adjusted for mean-centered age, mean-centered mean framewise displacement, and handedness (see Figure 2; lateralization for all eight lateralized networks in the Complete dataset is depicted in Supplementary Figure S3). The symmetrical Language network in the autism group appears to be driven by increased surface area in the right hemisphere.

**Figure 2.**
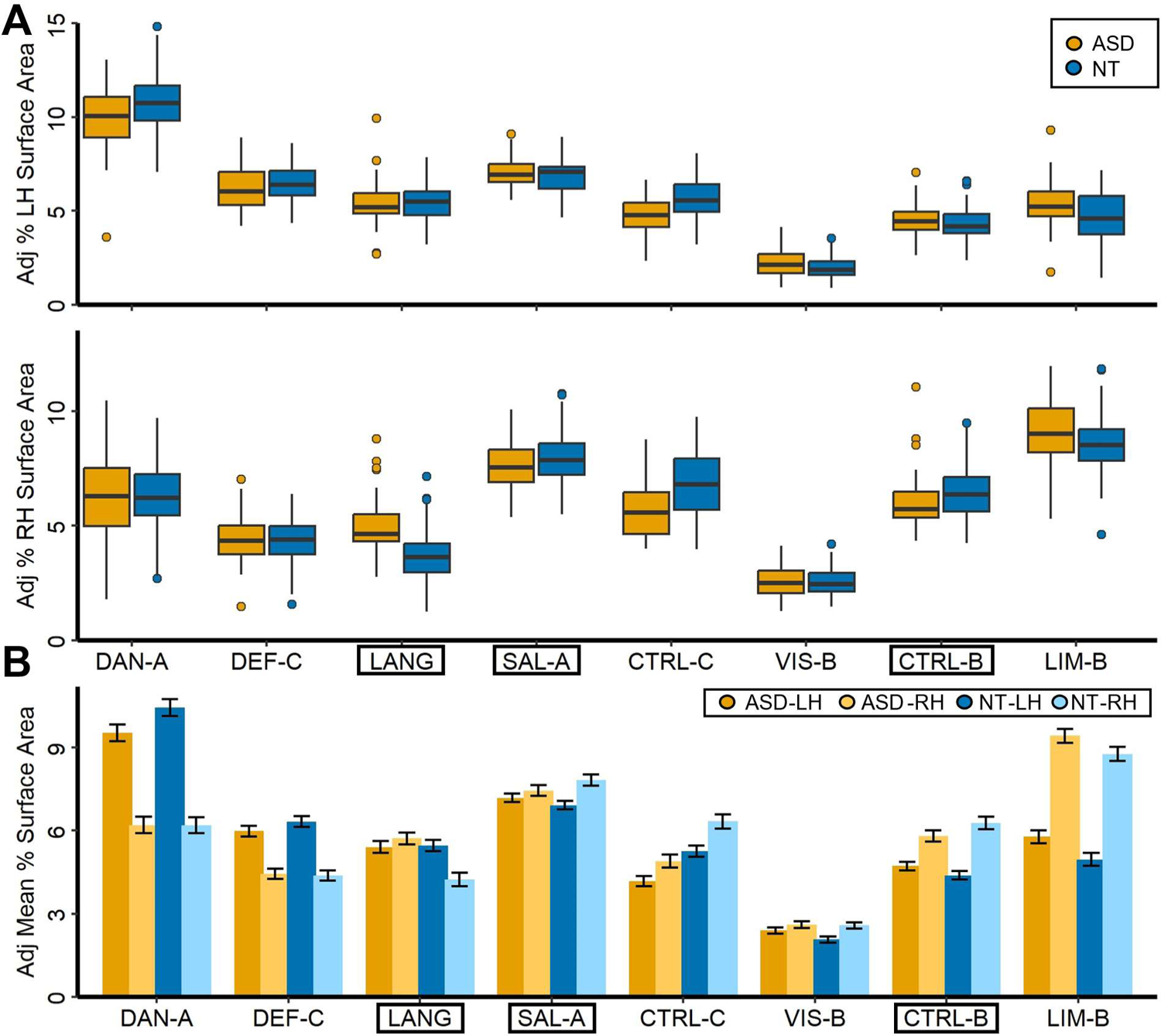
Percent surface area for 17 networks in ASD and NT individuals. Depicted in the top of Panel A is the model-adjusted percentage of the left hemisphere surface area occupied by a given lateralized network. Percent surface area was adjusted using the following formula: Surface area_adjusted_ = Surface area_raw_ - [β_1_(mean-centered age_raw_ - mean of mean-centered age_raw_) + β_2_(mean-centered FD_raw_ - mean of mean-centered FD_raw_) + β_3_(group_raw_ - mean group_raw_) + β_4_(handedness_raw_ - mean handedness_raw_)]. Depicted in the bottom portion of Panel A is the model-adjusted percentage of the right hemisphere surface area occupied by a given network. Points represent individual outliers. Depicted in Panel B is the adjusted mean percentage of surface area occupied by a lateralized network, with 95% confidence intervals. The left and right hemisphere estimates are displayed side-by-side. Black boxes have been used to indicate the networks for which a significant group difference was found.

Given the significant difference in mean framewise displacement between the ASD and NT groups, a sub-analysis of participants matched on mean framewise displacement (*N* = 96, the Complete dataset) was used. Similar conclusions to the unmatched analysis were reached, with group differences in lateralization identified for the Language (*t*(91) = −4.48, *p*-adjusted = .001), Salience/Ventral Attention-A (*t*(91) = 2.65, *p*-adjusted = .03), and Control-B (*t*(91) = 2.89, *p*-adjusted = .01) networks.

Previously, a significant group difference in the percent available volumes was identified, so a separate sub-analysis of participants matched on percent volumes available (*N* = 96, the Complete dataset) was undertaken. As with the unmatched analysis, group differences in lateralization were identified for the Language (*t*(91) = −4.35, *p*-adjusted < .001), Salience/Ventral Attention-A (*t*(91) = 2.69, *p*-adjusted = .02) networks, but not for the Control-B network (*t*(91) = 2.32, *p*-adjusted = .07).

Likewise, given the significant difference in full-scale IQ between the ASD and NT groups, a separate sub-analysis of participants matched on full-scale IQ scores was undertaken (*N* = 84, the Complete dataset). Group differences in lateralization was identified for the Language (*t*(79) = −3.71, *p*-adjusted = .001), Salience/Ventral Attention-A (*t*(79) = 2.67, *p*-adjusted = .03), and Control-B (*t*(79) = 3.21, *p*-adjusted = .01) networks.

### Verbal Ability, ASD Symptom Severity and Language Lateralization

To examine the potential relationship between verbal ability (measured via verbal IQ and Language network lateralization, a multiple regression with the covariates of mean-centered age, mean-centered mean framewise displacement, and handedness was used (*N* = 87; ASD = 45, NT = 42). Language lateralization was not a significant predictor of verbal IQ (*t*(81) = −0.63, *p* = .53).

Next, the relationship between language lateralization and autism symptom severity (measured via ADOS CSS scores, *N* = 47 ASD) was examined using a multiple regression with the covariates of mean-centered age, mean-centered mean framewise displacement, and handedness. Language lateralization was not a significant predictor of ADOS CSS scores (*t*(42) = 1.1, *p* = .28).

### Language Delay and Language Lateralization

The potential relationship between language delay and language lateralization was investigated using a multiple regression with the covariates of mean-centered age, mean-centered mean framewise displacement, and handedness. A significant group difference was found between the ASD with Language Delay and NT groups (*t*(109) = 4.62, *p* < .001, Cohen’s *d* = 1.05; see Figure 3). A significant group difference was also found between the ASD No Language Delay and NT groups (*t*(109) = −2.44, *p* = .02; Cohen’s *d* = 0.69). No significant group difference between the ASD Language Delay and ASD No Language Delay groups was found (*t*(109) = 1.21, *p* = .23).

**Figure 3.**
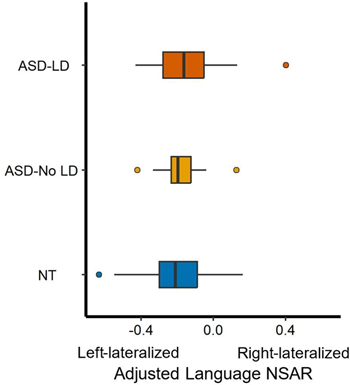
Language lateralization and language delay. Participants were binned into NT (*N* = 70), ASD Language Delay (*N* = 29), and ASD No Language Delay (*N* = 16), with three participants missing language delay data. On the y-axis are model-adjusted NSAR values for the Language network, with negative values representing left hemisphere lateralization and positive values representing right hemisphere lateralization. NSAR values were adjusted by regressing out the effects of mean-centered age, mean-centered mean framewise displacement, and handedness using the following formula: NSAR_adjusted_ = NSAR_raw_ - [β_1_(mean-centered age_raw_ - mean of mean-centered age_raw_) + β_2_(mean-centered FD_raw_ - mean of mean-centered FD_raw_) + β_3_(group_raw_ - mean group_raw_) + β_4_(handedness_raw_ - mean handedness_raw_)]. NSAR adjustment occurred separately for each group. A significant group effect on language lateralization was found between the NT and ASD Language Delay groups (*t*(109) = 4.62, *p* < .001) and between the NT and ASD No Language Delay groups (*t*(109) = −2.44, *p* = .02). Circles represent group mean adjusted NSAR values while bars represent the standard error of the mean.

## Discussion

In this study, we examined network lateralization in autistic and neurotypical individuals using a network surface area-based approach. We first hypothesized that group differences in lateralization would be constrained to areas associated with language. As expected, we identified differences in lateralization for the Language network. However, differences in network lateralization did not end there, and included the Salience/Ventral Attention-A and Control-B networks. Common among these three group differences was a reduction in asymmetry in the autism group, trending towards symmetric distributions. Additionally, of these three networks, the group difference in lateralization for the Language network showed the greatest effect size. Together, these findings evidenced a nuanced pattern of differences in network lateralization in autism, which were not restricted to the Language network, nor were they pervasive across all examined lateralized networks.

Next, we explored the connection between behavioral phenotypes and language lateralization. No significant relationships between verbal ability or autism symptom severity and language lateralization were found. However, language delay was identified as a stratification marker of language lateralization, with the greatest group difference found between the ASD with Language Delay and NT groups. This result suggests that the difference in language lateralization between the ASD and NT groups was predominantly driven by autistic individuals who experienced delayed language onset during development and does not reflect current language ability. In combination with prior research, this also suggests that differences in language lateralization occurring early in development are responsible for the differences in language lateralization in autism observed in the present study. Taken as a whole, these results provide further evidence for differences in functional lateralization in ASD, which appear to be behaviorally and clinically relevant in the case of language lateralization and language delay.

In the context of our research, network lateralization refers to the organizational principle whereby specific brain networks are predominantly based in one hemisphere versus distributed equally between both hemispheres. Of particular significance to our investigation is the lateralization of brain regions associated with language, given that language dysfunction is addressed within both categories of diagnostic criteria for the autism diagnosis. Our exploration extended beyond examining lateralization within eight networks, including the Language network, previously identified as lateralized in neurotypical individuals (59). We also sought to uncover relationships between language lateralization and three behavioral phenotypes, which together with prior studies, point to a developmental timeframe in which differences in language lateralization in autism emerge.

### Evidence for Differences in Functional Lateralization in ASD

The present study shed light on three networks—Language, Salience/Ventral Attention-A, and Control-B—where lateralization differed between ASD and NT individuals. Previously, language regions have been implicated in connectivity and asymmetry differences, leading to the postulation of the left hemisphere dysfunction theory of autism (13). Interestingly, the direction of group differences identified here indicates that the Language network in ASD is less asymmetrical than in NT individuals (see Figure 1 Panels B-D). This appears to be driven by an increase in Language network surface area in the right hemisphere compared with the NT group (see Figure 2). Other functional work has similarly identified a rightward shift in asymmetry in autism (14–18).

Perhaps, as suggested by the expansion-fractionation-specialization hypothesis, differences in the fractionation or specialization of the interdigitated theory of mind and language networks may contribute to the development of autism symptoms (63). This hypothesis proposes that as the cerebral cortex expands, certain core organizing areas act as anchors, while areas farther from these anchors self-organize into association cortex (64). These untethered association regions may exhibit a proto-organization at birth, which then fractionates and specializes through processes including competition and inherent connectivity differences. Any differences in the processes of expansion or fractionation may impact network specialization and potentially network lateralization. Considering the interdigitated nature of functional networks such as language and default networks, disturbances in the expansion or fractionation of one core area are likely to impact multiple networks both directly and indirectly. Our findings, demonstrating differences in lateralization across multiple networks in autism, align with this hypothesis.

The present study also identified a decrease in lateralization in the Control-B network in ASD. A mapping between resting-state functional connectivity and task activation has identified an executive control network as being associated with action–inhibition, emotion, and perception–somesthesis–pain (65). This has since been disentangled into two functional distinct control networks, which are linked to initiating and adapting control and the stable maintenance of goal-directed behavior (66). In ASD specifically, prior evidence has supported differences in control network structure (67), as well as increased right-lateralization in frontoparietal network components (25). However, because there is no standardized network taxonomy (68), we cannot definitively determine if the previous findings in control and frontoparietal networks directly relate to the observed lateralization differences in the Control-B network in the present study.

Unexpectedly, our research revealed a decrease in lateralization within the Salience/Ventral Attention-A network in ASD compared with NT individuals. Although this outcome was surprising, it could be partly attributed to the individualized approach taken in the present study, which may be more sensitive to differences in lateralization than group-averaged approaches. Regardless, the salience network is thought to identify relevant stimuli from internal and external inputs in order to direct behavior and is distinct from executive control networks (69,70). Complementary in function to the salience network, the ventral attention network is involved in spatial selective attention (71,72). Our finding is intriguing considering that alterations in attention are among the most frequently reported cognitive deficits in ASD (73). Neuroimaging studies have also supported this observation. Of note, Farrant & Uddin (74) reported hyperconnectivity in the ventral and dorsal attention networks in children with ASD, while hypo-connectivity was observed in the dorsal attention network in adults. However, the present study specifically identified decreased lateralization in the ventral attention network in ASD. Regardless, a salience network dysfunction theory of ASD has been proposed, suggesting that deviations in the salience network and anterior insula in particular may contribute to social communication and theory of mind deficits in ASD (75,76).

### Language Delay as a Stratification Marker for ASD

The present study identified a significant difference in language lateralization between NT and ASD with Language Delay individuals, similar to a previous study which found that language delay explained the most variance in extreme rightward deviations of laterality in autism (28). This finding is notable considering the disparities in datasets and modeling techniques between this study and that of Floris et al. (28). In the prior study, gray matter voxels were the subject of laterality, as opposed to functional connectivity-derived language network surface area. Additionally, significant group differences were identified using individual deviations from a normative pattern of brain laterality across development rather than from group mean comparisons. Another challenge, highlighted by Marek et al. (77) and Liu et al. (78), is the difficulty of establishing relationships between scanner-derived data (such as functional connectivity) and out-of-scanner behavioral measures. This is of particular concern with the use of the ADI for determining language delay, since this measure is retrospective and susceptible to memory errors such as telescoping (79). Thus, there is a clear need for prospective investigations of the relationship between language delay and language lateralization.

Regardless of these challenges, the causal direction and origins of the relationship between language delay in ASD and language lateralization remain unknown. Bishop (80) proposed several explanations for these differences. It was suggested that genetic risk may lead to language impairment, subsequently resulting in weak laterality (the neuroplasticity model). Alternatively, genetic risk might independently cause weak laterality and language impairment (the pleiotropy model), or weak laterality caused by genetic risk could subsequently lead to language impairment (the endophenotype model). An alternative model suggested by Berretz and Packheiser (81) posits that within any given neurodevelopmental or psychiatric condition, there is a singular, distinct endophenotype uniquely associated with altered asymmetries. Evidence from Nielsen et al. (26) suggests that deficits in language development may result in the abnormal language lateralization observed in ASD. This is supported by several pieces of evidence observed in the present study as well as in unpublished data (59). Notably, no consistent age-related effects on lateralization were identified previously (59), and the present study evidenced no direct relationship between language lateralization and verbal ability. However, language delay was found to act as a stratification marker for language lateralization, with the greatest effect occurring between the ASD with Language Delay and NT groups.

Together, this suggests that differences in language lateralization occurring early in development (likely *in utero* or shortly after birth), could underlie the differences in language lateralization observed in autism.

### Limitations

It should be noted that the dataset chosen for this study has certain characteristics which restrict the generalizability of our findings. First, the participant sample consisted entirely of males, which restricts the applicability of our results to females with ASD and may overlook potential sex differences. Additionally, the overwhelming representation of high verbal and cognitive performance individuals within the dataset further impact the generalizability of our findings.

Further investigations should focus on replicating these findings in larger and more diverse samples, as well as exploring the longitudinal trajectories of network lateralization in individuals with ASD. Given the present evidence suggesting that differences in language lateralization may be occurring early in development, it may be informative to explore differences in network lateralization in infancy and early childhood. Additionally, the incorporation of multimodal neuroimaging techniques could provide a more comprehensive understanding of the relationship between language network lateralization and language delay in ASD.

## Conclusions

In this study, we examined network lateralization in ASD and NT individuals using an individual-level approach based on participant network parcellations. First, we hypothesized that group differences in lateralization would be constrained to language-relevant regions. We identified group differences in lateralization for the Language, Salience/Ventral Attention-A, and Control-B networks, evidencing a selective pattern of functional lateralization differences in autism rather than a pervasive one. Additionally, we hypothesized that language delay would stratify language lateralization, such that the greatest group differences would be found between the NT and ASD with Language Delay groups. Support for this hypothesis was found, suggesting that language lateralization is behaviorally and clinically relevant to autism.

## List of Abbreviations

ADI: Autism diagnostic interview

ADI-R: Autism diagnostic interview-revised

ADOS: Autism diagnostic observation schedule

ASD: Autism spectrum disorder

CBIG: Computational brain imaging group

CSS: Calibrated severity scores

fMRI: Functional magnetic resonance imaging

FSL: FMRIB software library

IQ: Intelligence quotient

MP2RAGE: Magnetization prepare 2 rapid acquisition gradient echoes

NIfTI: Neuroimaging informatics technology initiative

NSAR: Network surface area ratio

NT: Neurotypical

TE: Echo time

TR: Repetition time

## Declarations Ethics Approval and Consent to Participate

All data were obtained with assent and informed consent according to the University of Utah’s Institutional Review Board.

## Consent for Publication

Not applicable.

## Availability of Data and Materials

The data reported on in the present study can be accessed on the NIMH Data Archive under #2400 (https://nda.nih.gov/edit_collection.html?id=2400). Preprocessing and individual parcellation pipeline code are available through the CBIG repository on GitHub at https://github.com/ThomasYeoLab/CBIG. Code used to implement the processing pipelines and perform statistical analyses are also available on GitHub at https://github.com/Nielsen-Brain-and-Behavior-Lab/AutismHemisphericSpecialization2023.

## Author’s Contributions

**MP**: Conceptualization, Methodology, Software, Validation, Formal analysis, Investigation, Writing – original draft, Writing – review & editing, Visualization. **MBDP**: Methodology, Software, Formal analysis, Investigation, Resources, Data curation, Writing – review & editing, Project administration. **DLF**: Methodology, Writing – review & editing. **EDB**: Writing – review & editing, Project administration, Funding acquisition. **BZ**: Writing – review & editing, Project administration, Funding acquisition. **JBK**: Writing – review & editing, Project administration. **NL**: Methodology, Writing – review & editing, Project administration, Funding acquisition. **ALA**: Writing – review & editing, Project administration, Funding acquisition. **JEL**: Resources, Writing – review & editing, Project administration, Funding acquisition. **JAN**: Conceptualization, Methodology, Investigation, Resources, Data curation, Writing – review & editing, Supervision, project administration. All authors read and approved the final manuscript.

## Acknowledgements

We thank former members of the Utah Autism CPEA for their assistance during the early stages of this project. We sincerely thank the children, adolescents, and adults with autism and the individuals with typical development who participated in this study, and their families. We are also grateful to Ru Kong for her assistance implementing the individual parcellation pipeline. Furthermore, we acknowledge the support of the Office of Research Computing at Brigham Young University.

