## Supplemental Figures and Tables for "Reduced Lateralization of Multiple Functional Brain Networks in Autistic Males"

Supplementary Materials

**Supplementary Table 1**

*Within-Dataset Replication Demographics*

|  | **Autism, *N =* 37** | | **Neurotypical, *N* = 60** | | **Group Comparison** | |
| --- | --- | --- | --- | --- | --- | --- |
|  | Mean (SD) | Range | Mean (SD) | Range | *t* | *p* |
| Age at Time 5 Scan (Years) | 27.93 (7.74) | 17.5 – 46.42 | 27.04 (6.72) | 16.33 – 44.42 | 0.57 | .57 |
| Percent Volumes Available | 82.34 (12.86) | 50.68 – 98.98 | 86.72 (10.33) | 55.72 – 100 | -1.75 | .08 |
| Handedness | 59.99 (47.37) | -82.15 – 100 | 73.61 (40.18) | -64.71 – 100 | -1.45 | .15 |
| Mean Performance IQ^a^ | 103.47 (16.44) | 67 – 134 | 116.31 (15.79) | 79 – 155 | -3.38 | .001 |
| Mean Verbal IQ^b^ | 101.5 (19.49) | 61 – 140 | 118.03 (10.78) | 99 – 140 | -4.45 | < .001 |
| Mean Full-scale IQ^c^ | 102.75 (17.79) | 60 – 137 | 118.78 (11.74) | 90 – 141 | -4.51 | < .001 |
| ADOS CSS^d^ | 7.78 (1.97) | 2 – 10 | - | - | - | - |
| ADI-R^e^ | 28.89 (6.73) | 14 – 40 | - | - | - | - |

^a^Mean Performance IQ: Autism *N* = 36, Neurotypical *N* = 36.

^b^Mean Verbal IQ: Autism *N* = 36, Neurotypical *N* = 36.

^c^Full-scale IQ: Autism *N* = 36	, Neurotypical *N* = 36.

^d^ADOS CSS at Study Entry *N* = 31; ADOS CSS scores at wave 5 *N* = 5.

^e^ADI-R: Autism *N* = 35.

**Supplementary Table 2**

*Within-Dataset Replication Data Quality (Mean Framewise Displacement)*

| **Group** | **Dataset** | **Mean (SD)** | **Range** | ***t*** | ***p*** |
| --- | --- | --- | --- | --- | --- |
| Autism | Discovery | 0.08 (0.03) | 0.04 – 0.19 | - | - |
| Neurotypical | Discovery | 0.07 (0.02) | 0.04 – 0.17 | - | - |
| Autism | Replication | 0.08 (0.02) | 0.05 – 0.13 | - | - |
| Neurotypical | Replication | 0.08 (0.03) | 0.04 – 0.18 | - | - |
| Autism vs Neurotypical | Discovery | - | - | 1.74 | .09 |
| Autism vs Neurotypical | Replication | - | - | 1.43 | .16 |
| Autism | Discovery vs Replication | - | - | -0.07 | .95 |
| Neurotypical | Discovery vs Replication | - | - | -0.72 | .47 |

**Supplementary Table 3**

*Identifying Lateralized Networks Using Multiple Regressions in Neurotypical Individuals (N = 70) from the Complete Dataset*

| Network Intercept | β | *SE* | *t* | *p*-adjusted |
| --- | --- | --- | --- | --- |
| Visual-A | .01 | 0.01 | 1.96 | .85 |
| **Visual-B** | 0.11 | 0.03 | 3.69 | .007 |
| Somatomotor-A | 0.01 | 0.02 | 0.47 | 10.93 |
| Somatomotor-B | -0.02 | 0.01 | -1.09 | 4.75 |
| **Language** | -0.19 | 0.05 | -4.11 | .002 |
| **Dorsal Attention-A** | -0.27 | 0.04 | -7.79 | < .001 |
| Dorsal Attention-B | 0.01 | 0.03 | 0.48 | 10.76 |
| **Salience/VenAttn-A** | 0.05 | 0.02 | 3.12 | .05 |
| Salience/VenAttn-B | -0.02 | 0.02 | -0.87 | 6.56 |
| Control-A | 0.01 | 0.01 | 0.52 | 10.32 |
| **Control-B** | 0.19 | 0.03 | 6.18 | < .001 |
| **Control-C** | 0.06 | 0.02 | 3.15 | .04 |
| Default-A | 0.02 | 0.03 | 0.79 | 7.37 |
| Default-B | 0.01 | 0.01 | 0.51 | 10.39 |
| **Default-C** | -0.2 | 0.03 | -6.04 | < .001 |
| Limbic-A | -0.01 | 0.02 | -0.53 | 10.15 |
| **Limbic-B** | 0.28 | 0.04 | 6.47 | < .001 |

*Note:* Coefficients and *p*-values for the intercept are shown. Networks with significant intercepts following Bonferroni correction are bolded.

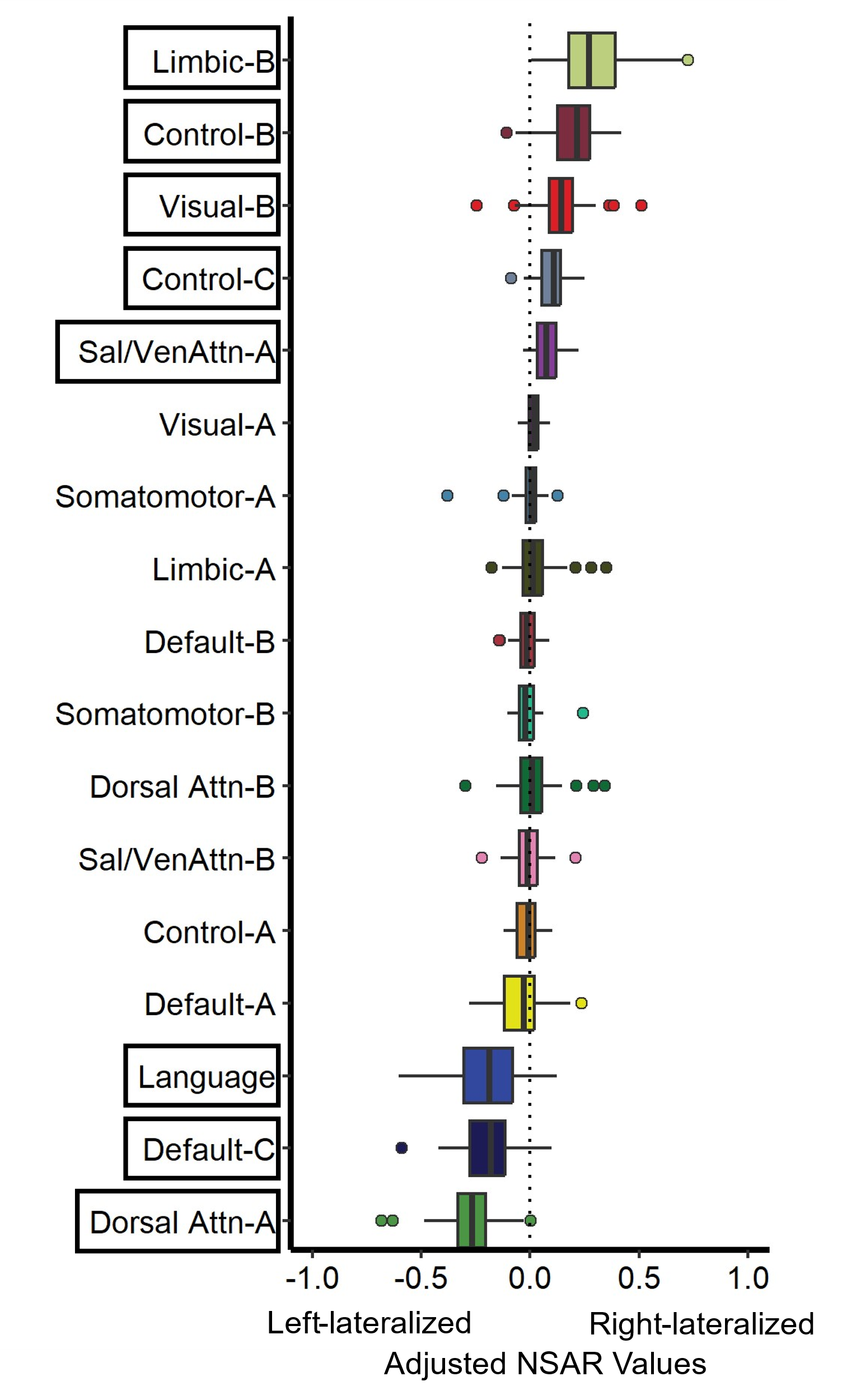
**Figure S1.** Lateralized networks in NT individuals from the Complete dataset. On the y-axis are the 17 networks and on the x-axis are model-adjusted NSAR values, with negative values representing left hemisphere lateralization and positive values representing right hemisphere lateralization. NSAR values were adjusted by regressing out the effects of mean-centered age, mean-centered mean framewise displacement, and handedness using the following formula: NSAR_adjusted_ = NSAR_raw_ - [β_1_(mean-centered age_raw_ - mean of mean-centered age_raw_) + β_2_(mean-centered FD_raw_ - mean of mean-centered FD_raw_) + β_3_(handedness_raw_ - mean handedness_raw_)]. NSAR adjustment occurred separately for each network. Eight networks were found to be significantly lateralized: Visual-B, Language, Dorsal Attention-A, Salience/Ventral Attention-A, Control-B, Control-C, Default-C, and Limbic-B. Black boxes have been used to indicate the lateralized networks.

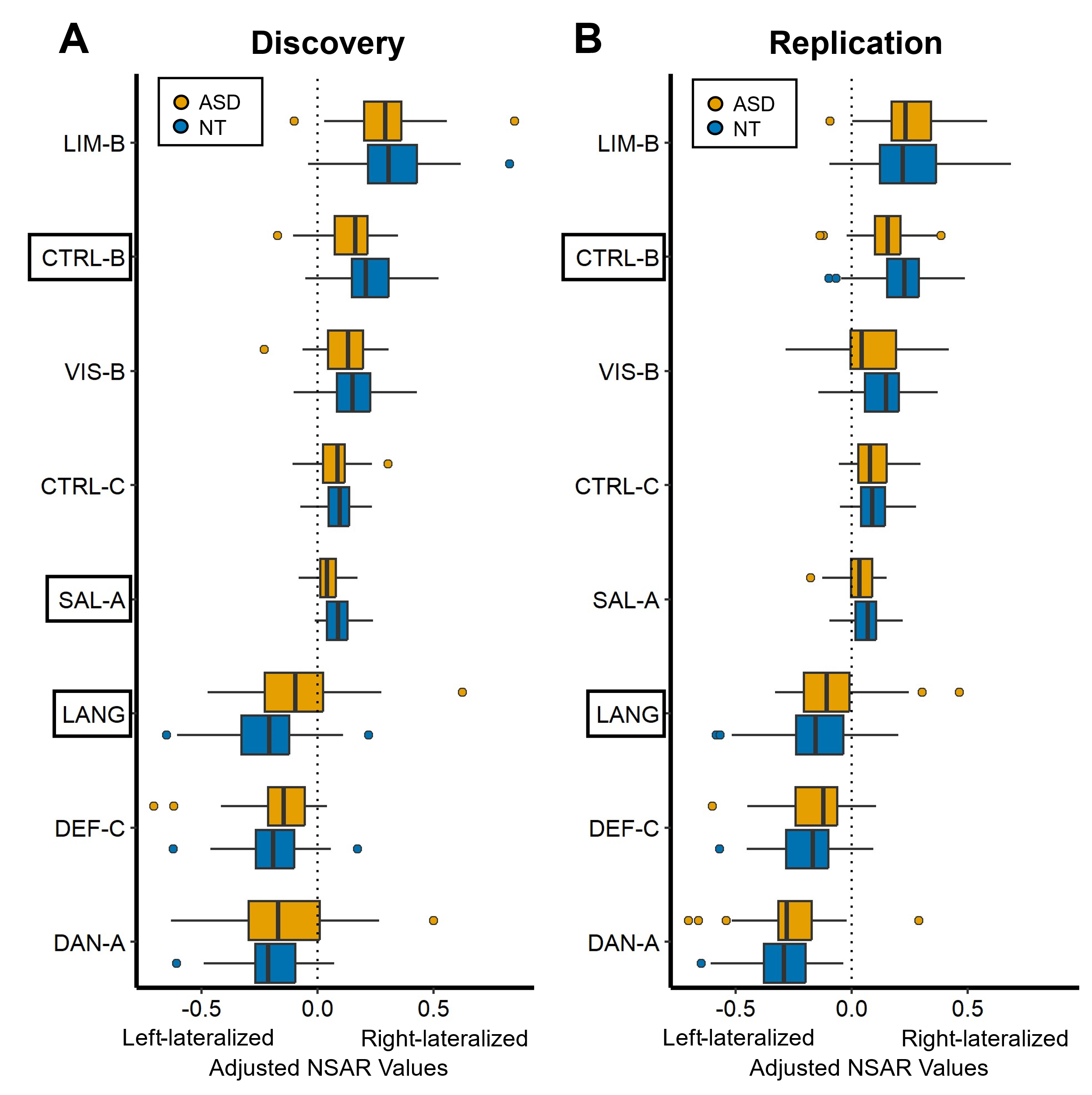
**Figure S2.** Lateralization across the Discovery and Replication datasets in autistic and neurotypical individuals. On the y-axis are the eight lateralized networks and on the x-axis are model-adjusted NSAR values, with negative values representing left hemisphere lateralization and positive values representing right hemisphere lateralization. NSAR values were adjusted by regressing out the effects of mean-centered age, mean-centered mean framewise displacement, and handedness using the following formula: NSAR_adjusted_ = NSAR_raw_ - [β_1_(mean-centered age_raw_ - mean of mean-centered age_raw_) + β_2_(mean-centered FD_raw_ - mean of mean-centered FD_raw_) + β_3_(group_raw_ - mean group_raw_) + β_4_(handedness_raw_ - mean handedness_raw_)]. NSAR adjustment occurred separately for each network and each group. A significant group effect on lateralization was found for three networks following Bonferroni correction in the Discovery dataset: Language (*t*(92) = -3.18, *p*-adjusted = .02), Salience/Ventral Attention-A (*t*(92) = 3.82, *p*-adjusted = .002), and Control-B (*t*(92) = 3.06, *p*-adjusted = .02). Significant group differences in lateralization for the Language (*t*(92) = -2.44, *p*-adjusted = .05) and Control-B (*t*(92) = 2.55, *p*-adjusted = .04) networks were replicated in the Replication dataset. Black boxes have been used to indicate the networks for which a significant group difference was found.

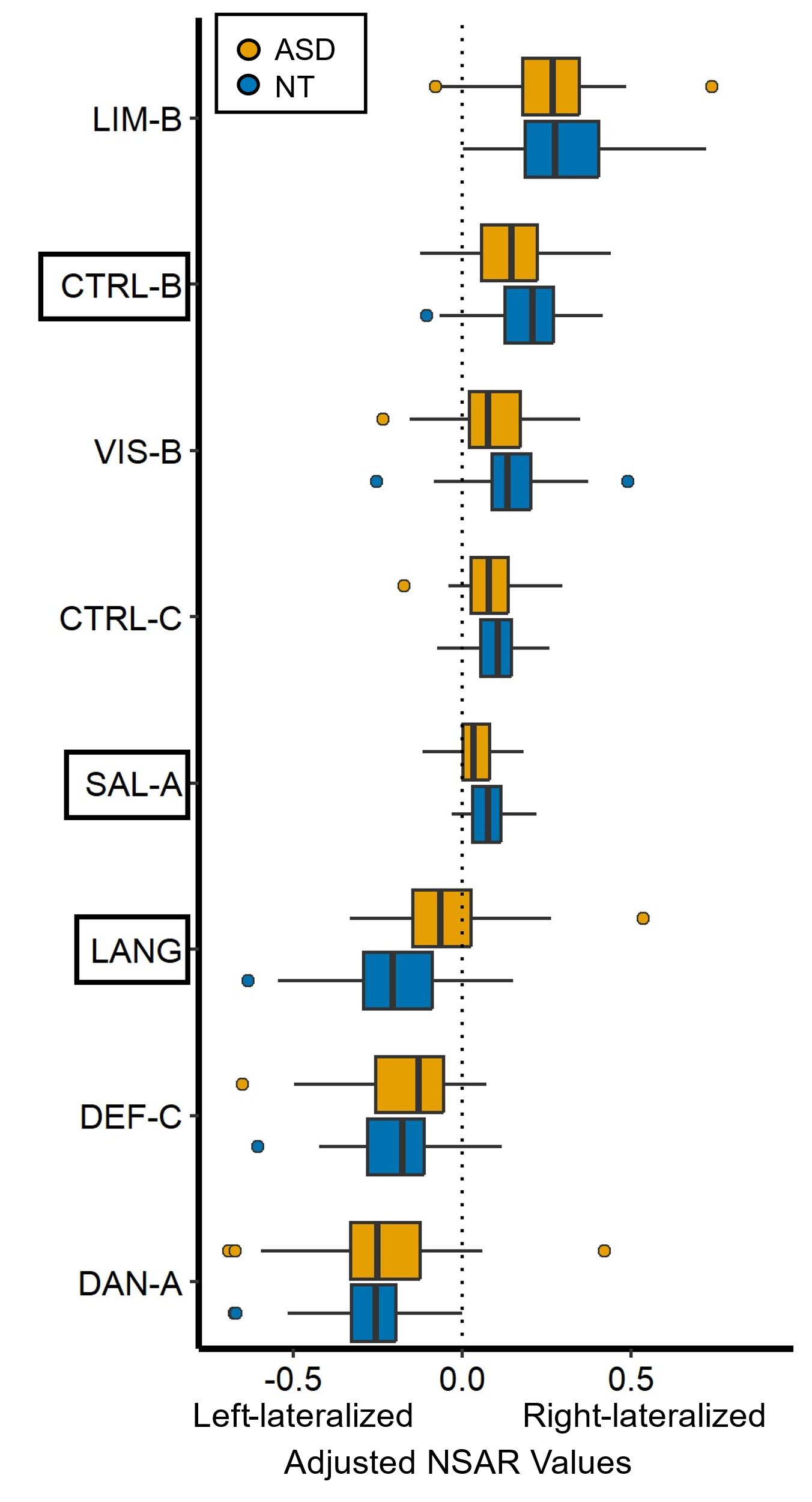
**Figure S3.** Lateralization for eight networks in the Complete dataset. On the y-axis are the eight lateralized networks and on the x-axis are model-adjusted NSAR values, with negative values representing left hemisphere lateralization and positive values representing right hemisphere lateralization. NSAR values were adjusted by regressing out the effects of mean-centered age, mean-centered mean framewise displacement, and handedness using the following formula: NSAR_adjusted_ = NSAR_raw_ - [β_1_(mean-centered age_raw_ - mean of mean-centered age_raw_) + β_2_(mean-centered FD_raw_ - mean of mean-centered FD_raw_) + β_3_(group_raw_ - mean group_raw_) + β_4_(handedness_raw_ - mean handedness_raw_)]. NSAR adjustment occurred separately for each network and each group. A significant group effect on lateralization was found for three networks following Bonferroni correction: Language (*t*(113) = -4.69, *p*-adjusted < .001), Salience/Ventral Attention-A (*t*(113) = 2.89, *p*-adjusted = .01), and Control-B (*t*(113) = 2.71, *p*-adjusted = .02). Black boxes have been used to indicate the networks for which a significant group difference was found.
